# Microsaccades track shifting but not necessarily maintaining covert visual-spatial attention

**DOI:** 10.1101/2025.07.14.664740

**Authors:** Anna M. van Harmelen, Freek van Ede

## Abstract

Covert attention enables the brain to prioritise relevant visual information without directly looking at it. Ample studies have linked covert visual-spatial attention to the direction of small fixational eye movements called microsaccades, offering researchers and clinicians a potential non-invasive tool to track internal states of covert attention. However, other studies have reported only a weak link or no link at all. We now show that the link between covert visual-spatial attention and microsaccades critically depends on the stage of attentional deployment. Across two experiments—each employing a distinct but widely adopted approach to fixational control—we show that spatial microsaccade biases were more pronounced (Experiment 1) or even exclusively present (Experiment 2) during the initial stage of shifting covert spatial attention, even when covert attention was subsequently maintained as testified by enhanced visual discrimination. This shows that the involvement of the brain’s oculomotor system in covert visual-spatial attention qualitatively changes over the course of attentional deployment, and how its peripheral fingerprint in the form of microsaccades reliably indexes shifting but not necessarily maintaining covert visual-spatial attention.

**Significance statement:** Covert attention is essential for efficient processing of information from our surroundings. Several studies have linked covert attention to the direction of miniscule eye-movements known as microsaccades. Other recent studies have questioned this link. We now explain this discrepancy by treating ‘attention’ not as a single static construct but as a dynamic process. We show that the link between microsaccades and covert attention critically depends on the stage of attention—with microsaccades robustly tracking shifting but not maintaining covert attention. This reveals how the contribution of the brain’s oculomotor system to attention changes dynamically over time. These findings have implications for the use of microsaccades as an accessible marker of attentional (mal)function across psychological and neurological conditions.

## Introduction

Visual-spatial attention refers to the dynamic process by which the human brain selects and prioritises the processing of currently relevant visual information (1–3). Visual-spatial attention can operate in two modes: overtly, by physically directing gaze towards relevant stimuli, and covertly, by attending to peripheral visual information outside of the current fixation. Extensive research has established that the neural circuitry that mediates shifts of overt visual-spatial attention, namely the oculomotor system, also contributes to covert visual-spatial attention (4–8). A canonical behavioural manifestation of this overlap in neural circuitry comes from the study of microsaccades—small eye movements that occur even during attempted fixation (9–13). In particular, many studies have reported an association between spatial biases in microsaccade direction and the allocation of covert visual-spatial attention.

This link between microsaccades and covert visual-spatial attention has been demonstrated repeatedly. Early studies linked the direction of microsaccades to the deployment of covert attention (14, 15) and these findings were later replicated and extended. For example, it has been demonstrated in both humans (14–25) and non-human primates (26, 27); during both externally directed perceptual attention (14, 15, 17, 20, 21, 24–27) and internally directed attention within visual working memory (16, 18, 19, 22, 23); and in both perception and action tasks following directional cues (28). Several studies have further linked the directional microsaccade bias to task performance (18, 21, 25, 28, 29). For example, following spontaneous microsaccades, perception of visual targets presented in the same direction is better (25), and visual discrimination benefits may start already prior to microsaccade execution (21). Recent evidence further suggests that microsaccades may even play a causal role in shaping the perception of peripheral stimuli (30).

At the same time, the link between covert visual-spatial attention and microsaccades is not always found (31–33), leading to mixed results in the literature. Notably, in a recent article, Willett & Mayo (33) reported no bias in microsaccade directions, despite clear behavioural and neural demonstrations that covert attention was allocated to the relevant visual stimulus.

One possible source underlying the mixed results that pervade the literature may be that ‘attention’ has been treated as a unitary and static construct. This is problematic under the hypothesis that the link between microsaccades and covert attention may only be reliable for specific stages associated with the deployment of spatial attention, such as when shifting between locations, but not when subsequently maintaining attention at the new location (34). Indeed, while most studies that reported a link used tasks that required *shifting* covert visual-spatial attention, the recent study by Willet & Mayo (33) that questioned this link used a task that required *maintaining* covert visual-spatial attention after it had already been shifted. Thus, whether—and how reliably—microsaccades track covert visual-spatial attention may fundamentally depend on whether one considers the period of shifting attention, or the ensuing period of maintaining attention after the shift (34).

This differentiation by stage of attention also naturally follows from the putative function of the brain’s oculomotor system in attention. Consider the case of overt attention: when aiming to attend to an object by looking at it, the oculomotor system is initially engaged once to shift the eyes. Once the eyes have landed on the attended location, no further eye-movements are necessary, yet overt attention can remain at this location. We pose that the same may apply for covert attention: shifting covert attention may engage the oculomotor system particularly to relocate the focus of covert attention, leading to a bias in microsaccade direction particularly while the shift occurs. Once shifted, visual-spatial attention may effectively remain at the covertly attended location, but with a distinct contribution from the oculomotor system that does not necessarily result in a bias of microsaccade direction.

To address the hypothesis (34) that microsaccades may particularly track *shifting* covert visual-spatial attention—in comparison to *maintaining* attention at this location after the initial shift—we conducted two complementary experiments that each employed a distinct but widely adopted approach to gaze control during the task. Consistent with our hypothesis, microsaccades predominantly (Experiment 1) or exclusively (Experiment 2) tracked shifting compared to maintaining covert visual-spatial attention, even though covert attention was maintained at the attended location after the initial shift, as evidenced by visual discrimination performance.

## Results

Forty-eight healthy human volunteers performed a covert visual-spatial attention task in which a central colour cue indicated which of two laterally presented stimuli (positioned on the left and right below central fixation) would be most likely to show the target event: a slight orientation change that had to be judged as being clockwise or anticlockwise (**Fig. 1A**). To be able to study covert attention across both shifting and subsequent maintaining phases, we varied the interval between cue and target from 500 to 3200 ms. This necessitated shifting attention immediately in response to the cue (as the target event could already occur 500 ms after cue onset) as well as maintaining it on the cued stimulus (since the target event would often occur after longer cue-target intervals). We conducted two versions of this task, differing only in the approach to fixational control during the task. In Experiment 1, central fixation was instructed and monitored by the experimenter. Experiment 2 included gaze-contingent trial termination whenever gaze deviated more than 2 degrees visual angle from central fixation, as controlled by custom software. Adopting both approaches enabled us to relate our findings to a broader set of prior studies given that both are widely used in the vast literature on covert visual-spatial attention.

**Figure 1.**
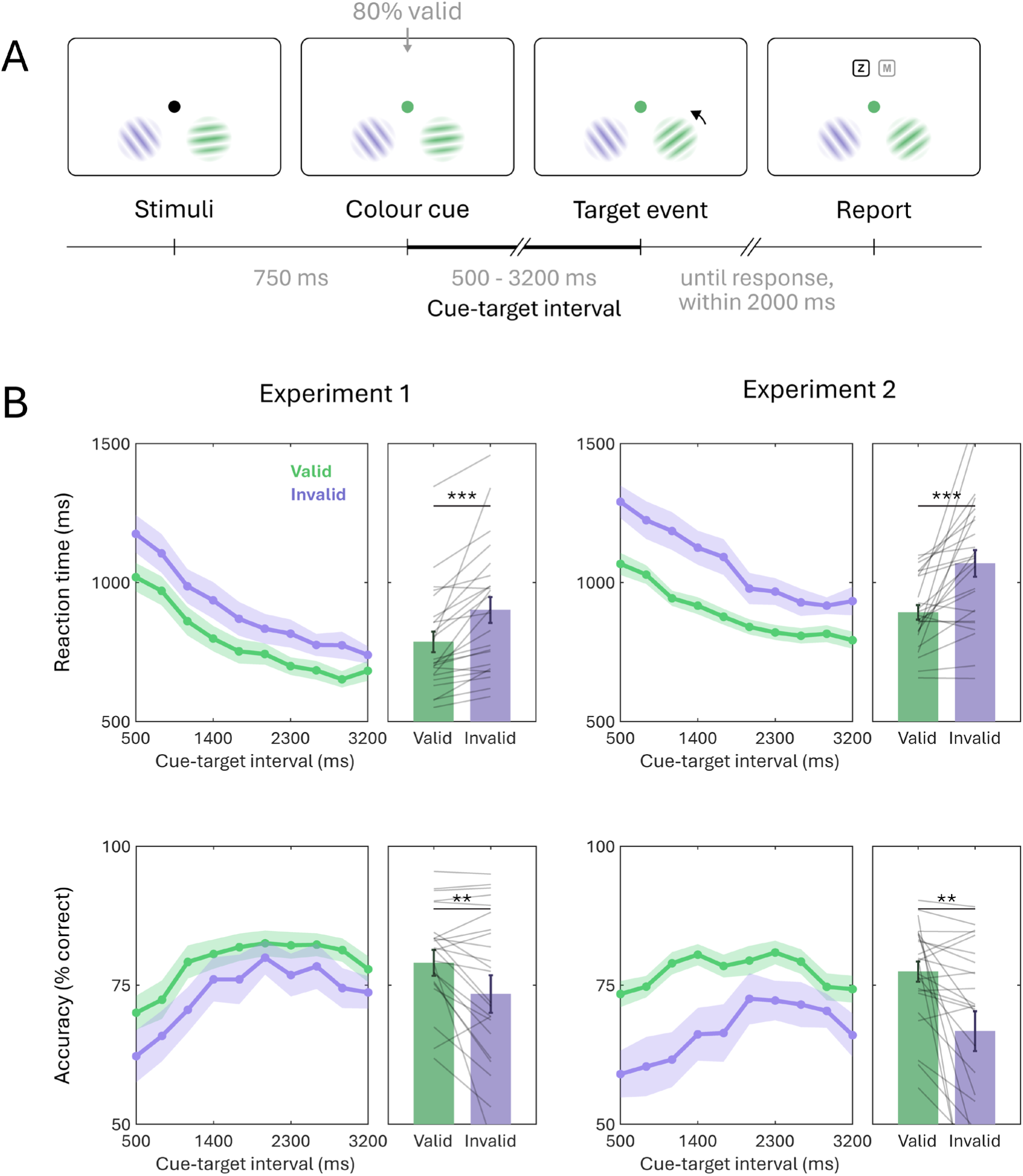
Attention was successfully shifted and maintained. (A) Schematic showing the progression of a single example trial. A colour change of the central fixation dot cued which of two visual stimuli was most likely to change orientation (the visual target). After a variable delay, the target event (the orientation change) occurred in either the cued (80% of trials, valid) or other (20%, invalid) stimulus and participants reported whether it was clockwise or anticlockwise. Colours shown serve only as an example, for the experimental colours, see the methods section. Supplementary Figure S1 shows the stimulus screen and stimulus sizes and distances to scale. (B) Behavioural performance (reaction times above, accuracy scores below) as function of cue-target interval for validly vs. invalidly cued targets. Circular markers indicate mean values, with shaded areas indicating the standard error of the mean. Bars reflect the mean behavioural performance, with whiskers indicating the standard error of the mean. Grey lines indicate individual participants’ data. Throughout the entire figure, the following significance levels were used: *: p< 0.05, **: p < 0.01, ***: p < 0.001, ****: p < 0.0001.

### Attention was successfully shifted and maintained

We first verified that our task was successful in inducing early shifts of visual-spatial attention that were maintained even for target events occurring several seconds after the cue that prompted the initial attention shift. As shown in **Figure 1B**, responses were faster and more accurate for target events occurring at the cued (valid) versus the other (invalid) stimulus, with the cue-related attentional modulation extending across all cue-target intervals. Cue-validity effects are summarised in the bar graphs in **Figure 1B**, and were robust in both experiments, for both reaction times (E1: t(23)=4.673, p=1.052e-04, d=0.540; E2: t(23)=4.098, p=4.407e-04, d=0.894) and accuracy (E1: t(23)= 3.281, p=0.003, d=0.383; E2: t(23)= 3.062, p=0.006, d=0.745). (For cue-validity effects at each of the tested cue-target intervals, see **Supplementary Table S1**.)

### Attention was accompanied by a bias in microsaccade direction

Building on prior demonstrations of a bias in microsaccade direction accompanying the spatial allocation of covert visual attention (see introduction), we assessed whether microsaccades are also biased towards the cued stimulus in our orientation change detection task. We were specifically interested in the *temporal dynamics* of such a microsaccade bias in our task, which invoked both shifting covert attention to the cued visual stimulus, as well as maintaining it there until the target event (the orientation change) would occur.

We exclusively analysed directional microsaccade biases prior to the orientation change, to ensure our results are uncontaminated from potential responses related to the target event itself. To ensure sufficient trials for analysing microsaccades, we included all trials where the orientation change occurred later than 1400 ms after cue onset. This provided a good trade-off between having sufficient trials, while also having sufficient time to capture microsaccade biases during the initial shifting phase and the ensuing phase in which attention was maintained on the cued stimulus after this shift. Importantly, our conclusions remain unaffected when investigating the entire time course through a complementary analysis (see **Supplementary Figure S2**).

Figure 2A shows the time-resolved rates of downwards saccades made either towards the cued stimulus or towards the other stimulus, for both experiments (for directional histograms and a visualisation of which saccades were included, see **Supplementary Figure S3**; a complementary analysis of upwards saccades can be found in **Supplementary Figure S4**). To zoom in on the ‘attentional modulation’, **Figure 2B** shows the spatial saccade bias, as the difference in saccade rates toward the cued versus the other stimulus. In both experiments, a significant saccade bias is present towards the cued item, starting around 200 ms after cue onset (E1: cluster p < 0.001; E2: cluster p=0.036). While in Experiment 1 saccade directions remain significantly biased towards the cued stimulus throughout the entire investigated period, in Experiment 2—where fixation demands were more strict, as trials were terminated upon breaking fixation—this bias became short lived, occurring only within the typical window associated with attentional shifting (see 14, 16, 18, 20, 23, 35–37). When directly comparing this spatial saccade bias between experiments, we observed that the effect was significantly larger in Experiment 1 than in Experiment 2 from 535 to 745 ms (cluster p=0.013) and 1153 to 1309 ms after cue onset (cluster p=0.029). A complementary analysis further showed that the magnitude of the spatial saccade bias correlated with the magnitude of the cue-related attentional modulation in Experiment 1, but not in Experiment 2 (see **Supplementary Figure S5**).

**Figure 2.**
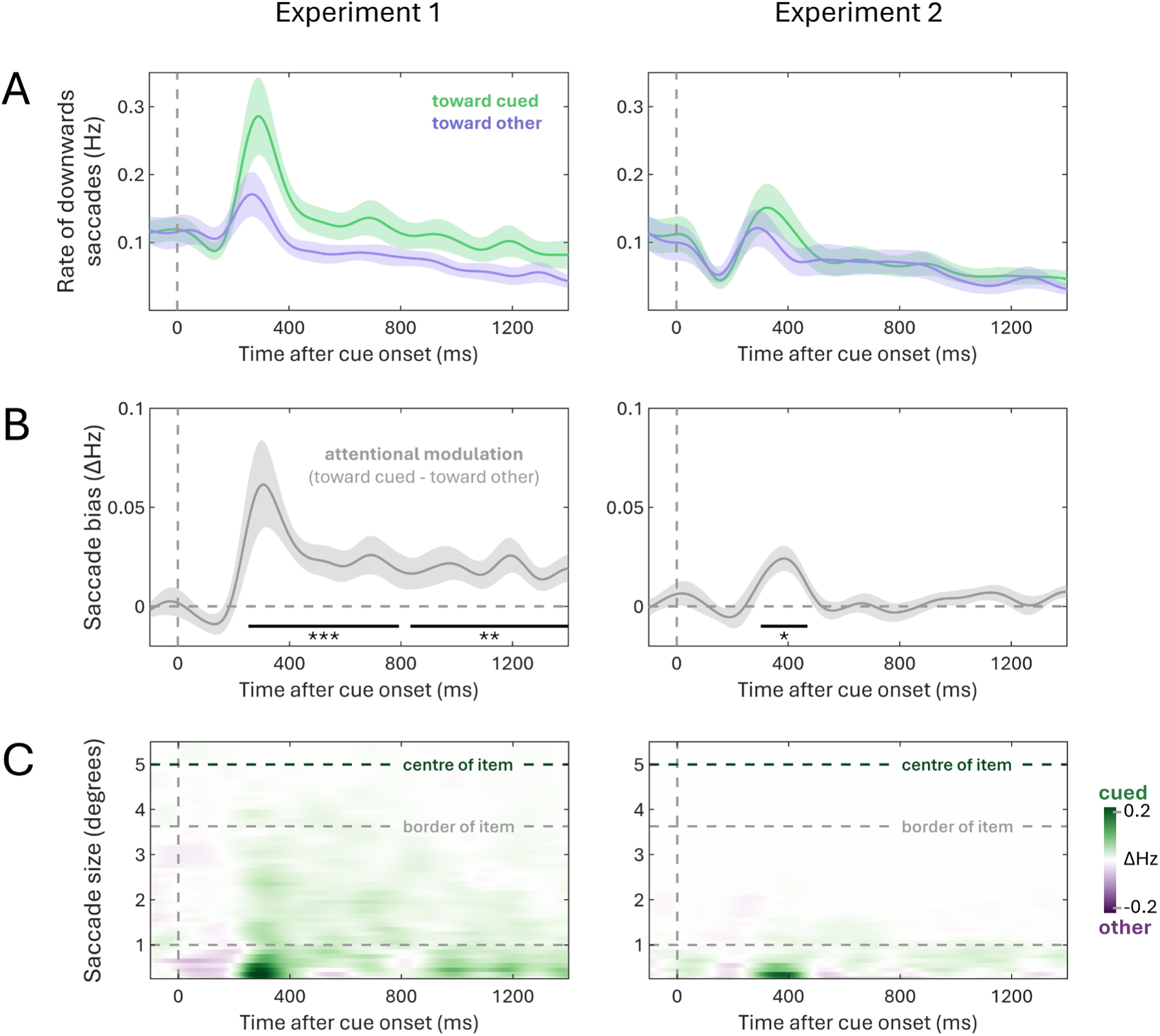
Attention was accompanied by a bias in microsaccade direction. (A) Time courses of saccade rates for saccades towards the cued and the other item. This analysis only includes downward saccades, as stimuli were positioned at a 45° angle below fixation, allowing us to distinguish saccades towards the item from those returning to central fixation. The time courses indicate mean values, with shaded areas indicating the standard error of the mean. (B) Time courses of the attentional modulation of saccade direction (toward cued – toward other). The time courses indicate mean values, with shaded areas indicating the standard error of the mean. Black horizontal lines indicate significant temporal clusters (as determined by a cluster-based permutation test). (C) Time courses of attentional modulation (as shown in B) as a function of saccade size. For reference, dashed horizontal lines indicate 1 degree visual angle, as well as the locations of the centre and closest border of the visual stimulus. Throughout the entire figure, the following significance levels were used: *: p< 0.05, **: p < 0.01, ***: p < 0.001, ****: p < 0.0001.

Rather than classifying saccades as ‘micro’ or ‘macro’ based on an inherently arbitrary threshold, **Figure 2C** shows the attentional modulation (saccade bias) as a function of the size of the saccades that contributed to it. Consistent with our instructions to deploy attention covertly, the reported bias was not driven by participants looking directly at the cued stimulus at 5 degrees visual angle from central fixation, but rather by smaller saccades that were *biased* in the direction of the cued stimulus. Indeed, the saccade bias was driven predominantly (Experiment 1) or exclusively (Experiment 2) by saccades in the classic microsaccade range, i.e. below 1 degree. Note that Figure 2C does not show saccade landing positions. While it is theoretically possible for multiple small unidirectional saccades to lead to a larger change in gaze position, a complementary analysis shows that fixation was maintained during the period of peak microsaccade rate in both experiments (see **Supplementary Figure S6**).

### Microsaccade directions are biased predominantly during attentional shifting

To summarise our findings, we finally quantified the observed spatial bias separately in two time windows that were chosen a-priori to capture either the period of ‘shifting’ or the ensuing period of ‘maintaining’ covert attention at the cued stimulus, after the initial shift. Specifically, we defined the shift period as the time period from 200 to 600 ms after cue onset, consistent with prior microsaccade studies (e.g., 14, 15, 18, 20, 23) as well as with complementary behavioural studies showing that voluntary attentional deployment unfolds between 200-600 ms after an endogenous attention cue (e.g., 35–37). Following the logic that participants were required to shift attention to the cued stimulus and retain it there, we defined the maintenance period simply as the interval immediately following the period of shifting: from 600 to 1400 ms after cue onset.

**Figure 3** shows a quantification of the directional bias in both timeframes. This confirmed a significant directional bias during the ‘shift’ period in both experiments (E1: t(23)=3.117, p=0.005, d=0.615; E2: t(23)= 2.543, p=0.018, d=0.502), while the bias in the ‘maintain’ period was significantly only in Experiment 1 (t(23)=2.838, p=0.009, d=0.560) and no longer present in Experiment 2 (t(23)=0.889, p=0.383, d=0.175). To clarify this result, we found moderate Bayesian evidence for the absence of a bias during the maintenance period in Experiment 2 (BF_01_=3.26). This corresponds with the absence of a significant cluster in Experiment 2 in **Figure 2**. Finally, complementing the findings for Experiment 1 shown in **Figure 2**—where we found a single cluster spanning both shift and maintenance periods—an additional targeted quantification revealed that in Experiment 1, the bias was still significantly larger during the shift than during the post-shift maintain period (t(23)=2.300, p=0.031, d=0.329).

**Figure 3.**
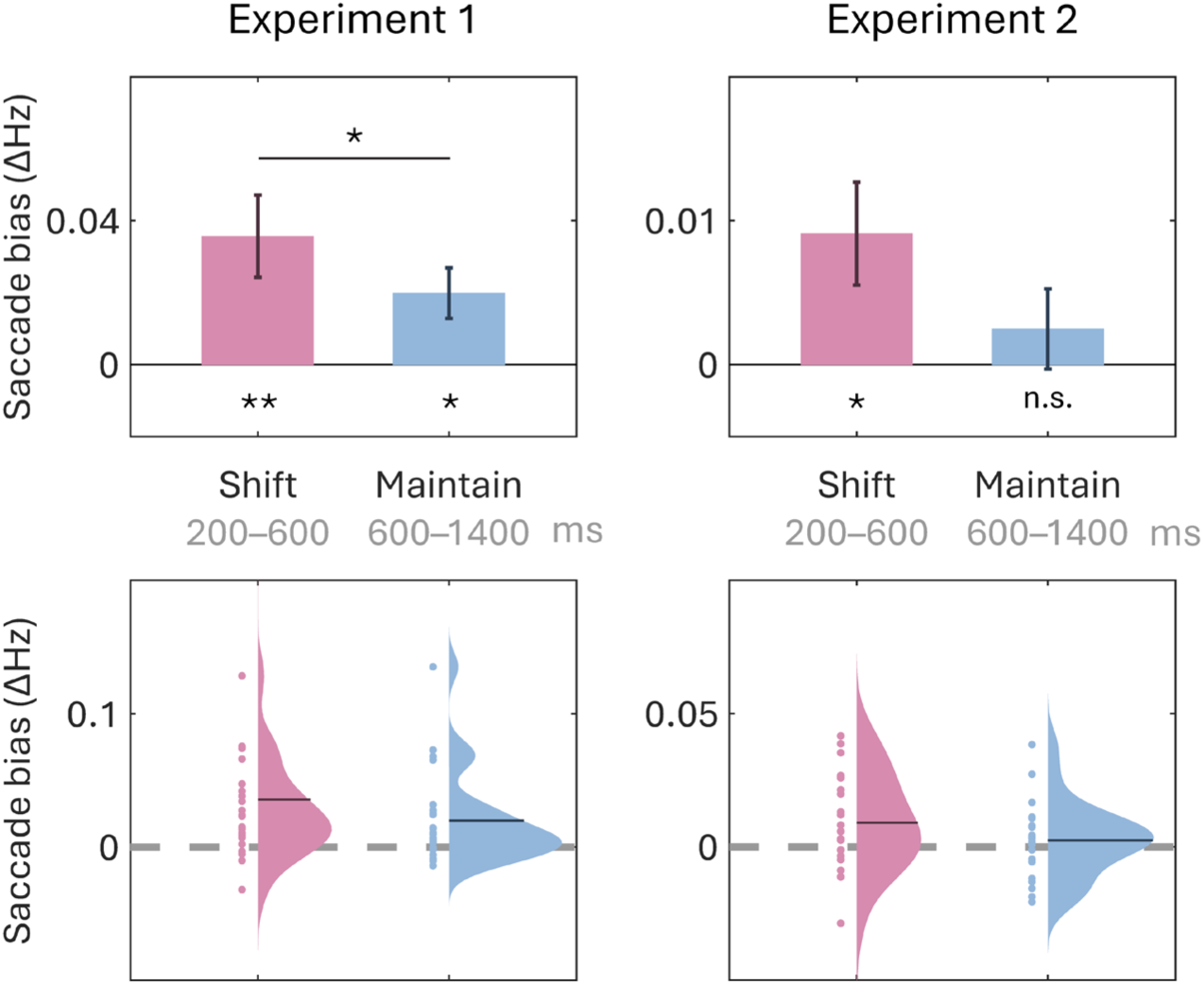
Microsaccade directions are biased predominantly during attentional shifting. Bars represent the mean saccade towardness rate in the denoted timeframes, with whiskers indicating the standard error of the mean. Distribution and scatter plots show individual participants’ data, with dark lines indicating the mean. Note that the y-axes have different scales between Experiment 1 and Experiment 2 and between bar graphs and distribution plots, for visualisation purposes. Throughout the entire figure, the following significance levels were used: *: p< 0.05, **: p < 0.01, ***: p < 0.001, ****: p < 0.0001.

## Discussion

By investigating attention as a dynamic process, we reveal how the relationship between microsaccades and covert visual-spatial attention critically depends on the stage of attentional deployment. Specifically, our results show that microsaccade directions are predominantly (Experiment 1) or even exclusively (Experiment 2) biased during the initial process of shifting covert attention to a relevant visual stimulus, as opposed to the subsequent stage of maintaining attention on this stimulus after the initial shift. We observed this even though visual discrimination performance confirmed the allocation of attention well beyond the initial shift window. The insight that the relation between microsaccade direction and covert visual-spatial attention depends on the stage of attention provides a critical nuance in our general understanding of the relationship between the oculomotor system and covert attention. This insight may not only clarify apparent inconsistencies in the literature (34), but also has important theoretical and practical implications, as we elaborate below.

When considering microsaccades as a peripheral marker of the oculomotor system (38, 39), our data suggest that role of the oculomotor system is likely distinct during different stages of covert attentional deployment. While the role of the wider oculomotor system in attention is well documented (4–8), the notion that this role may fundamentally differ across stages of attentional deployment (40) remains underappreciated. Yet, this dynamic stage-dependent participation is highly plausible when we consider that the same system is responsible for both covert and overt shifts of attention. In the case of overt attention, we make an eye movement to bring a new object into view once, and then simply keep this object foveated. By studying microsaccades, we provide evidence that a similar dynamic participation of the oculomotor system likely holds for covert visual-spatial attention. This does not necessarily imply that the oculomotor system contributes *more* during attentional shifts—it simply contributes differently. For example, shifting covert attention may principally recruit populations of motor and visual-motor neurons within the oculomotor system, while after attention has been shifted, covert attention may be held in place primarily by visually selective neurons in the same system that do not bias microsaccade directions (for the widely studied distinction between visual, motor, and visual-motor tuning, see e.g., 41, 42). Future work using direct neurophysiological recordings are required to asses this possibility.

Our findings also have practical implications. When using microsaccades as a marker to study covert attention, our data demonstrate that this marker is predominantly useful for tracking attentional shifts, but less so for tracking subsequent maintaining of visual-spatial attention (for at least one other recent study showing a transient effect likely confined to the attentional shift period, see 28). This does not necessarily negate its use to also track more sustained forms of attention (24, 43), but our data make clear how this is less reliable and may break down, especially when employing stricter fixational control (as was shown in Experiment 2). Furthermore, while our data confirm a robust link between microsaccade direction and shifts of covert visual-spatial attention on a trial average, it is important to note that microsaccades are not necessarily a reliable marker of attention shifts at the single-trial level. In fact, previous research has shown that microsaccades and attention shifts can decouple at the single-trial level (18, 31, 44, 45), and that attention shifts only influence the likelihood of microsaccades following the same direction as attention, without directly triggering any new microsaccades (19). Therefore, even if microsaccades may more reliably track shifting than maintaining attention, as our current findings show, our findings should not be taken as evidence that microsaccades reliably track attentional shifts at the single-trial level.

In this current work, we assessed the relationship between microsaccades and covert attention in two experimental settings with different approaches to fixation control: instructed fixation with monitoring by the experimenter (Experiment 1) or with gaze-contingent trial termination controlled by custom software (Experiment 2). Because both approaches are commonly employed in the field, this set-up allows our data to be comparable with a broader literature on covert attention. In both cases, microsaccades were more reliably associated with the initial shifting of attention. Moreover, during the stricter fixation demands of Experiment 2, the link between microsaccades and attention could exclusively be established during the initial shifting period. This was the case even though we observed clear modulations of performance at intervals far beyond the initial shift period. Together, these complementary experiments reinforce our central conclusion that the link between microsaccades and covert visual-spatial attention critically depends on the attention stage under investigation. In addition, they show how the exact mode of fixation control can shape the observed nature of this link.

The present study specifically examined microsaccade biases in relation to shifting and the ensuing period of maintaining covert visual-spatial attention in a task where the target event could occur anywhere between 500 and 3200 ms after cue onset. In future work, it will be valuable to extend our findings to situations where covert visual-spatial attention must be sustained over much longer timescales. Such situations may also naturally invoke lapses of attention that then require a re-establishing or re-shifting of covert visual-spatial attention. Whether such refocusing resembles the cue-induced shifts studied here—and is therefore similarly tracked by microsaccades—remains an interesting question for future research.

By considering attention not as a unitary and static construct but as a dynamic process, our data highlight how the involvement of the oculomotor system in covert visual-spatial attention fundamentally depends on the stage of attentional deployment, as reflected peripherally by microsaccades. This provides new insight into the intricate dynamics of covert visual-spatial attention and its oculomotor underpinnings and carries implications for the utility of microsaccades as an accessible peripheral marker of attention and its dysfunction in various neurological and psychiatric conditions.

## Methods

All experimental procedures were reviewed and approved by the ethics committee of the Vrije Universiteit Amsterdam. Each participant provided written informed consent prior to participation and was reimbursed with 10 euros/hour or participation credits.

### Participants

We performed two experiments that only differed in the approach to fixation control (detailed below under ‘*Experiment 1 vs. Experiment 2: Gaze-contingent trial termination*‘). A sample size of twenty-four per experiment was set a-priori, as based on previous publications from our lab with similar experimental designs and outcome measures.

All participants were recruited from the Vrije Universiteit Amsterdam. Twenty-five healthy human volunteers participated in experiment one, one of which was excluded post-hoc due to the inability to keep stable fixation. This resulted in the a-priori determined sample size of twenty-four participants (age range: 18-37; 19 women, 4 men and 1 non-binary; 21 right-handed). Twenty-nine healthy human volunteers participated in experiment two, five of which were unable to finish the experiment due to the inability to keep stable fixation. This resulted in the a-priori determined sample size of twenty-four participants (age range: 18-28; 20 women, 4 men; 23 right-handed). Recruitment for both experiments was done independently, with the constraint that participants were not allowed to participate in both experiments.

### Task and procedure

Experiment 1 and Experiment 2 used the same task that we depict in **Figure 1A**. This consisted of a covert visual-spatial attention task requiring participants to detect and discriminate a sudden orientation change in one of two presented Gabor patches. Prior to the orientation change, a colour cue was shown indicating with 80% reliability which Gabor patch would contain the target event (i.e., the orientation change).

Both experiments contained 20 blocks of 40 trials each. Before starting the experiment, participants practiced both the required response and entire trials until they felt acquainted with the task.

The stimuli would always be on the screen for 750 ms before the central fixation marker would change to match the colour of either stimulus serving as the cue. Left and right cues were equally likely and counterbalanced across trials. The interval between cue onset and the target event (the orientation change of either stimulus) varied from 500 to 3200 ms in discrete steps of 300 ms, resulting in ten possible cue-target intervals. Cue-target intervals were randomly drawn and counterbalanced across trials. The inclusion of early target events ensured that participants had an incentive to shift attention upon cue onset, while the inclusion of later target events enabled us to also study the period of subsequently maintaining attention on the cued stimulus.

After the orientation change occurred, participants reported the direction of the orientation change (i.e. clockwise or anticlockwise) by pressing either the ‘z’ (anticlockwise) or ‘m’ key (clockwise). Clockwise and anticlockwise target events were equally likely and counterbalanced across trials.

The Gabor patches were each positioned at an angle of 45° below the central fixation dot (see **Supplementary Figure S1** for a to-scale representation of the stimulus screen), one to the left and the other to the right. Critically, this positioning prevents conflating saccades associated with returning to fixation after moving in the direction of one stimulus with saccades directed towards the other stimulus.

### Experimental set-up

Both tasks were programmed in Python 3.8.17 using the PsychoPy library (46) to generate the stimuli. Participants were seated 70 cm in front of a 24-inch LCD monitor, with a 1920 x 1080 pixel resolution and a 239 Hz refresh rate.

Gabor patches presented on a dark grey background (rgb: 64, 64, 64, luminance: 29.0 cd/m^2^) could be one of four colours: blue (rgb: 19, 146, 206), magenta (rgb: 217, 103, 241), green (rgb: 101, 148, 14), orange (rgb: 238, 104, 60). Colours were randomly intermixed across trials with the constraint that the two Gabors in each trial had a different colour. All four colours were calibrated prior to the experiment to ensure equiluminance on the monitors used in the lab, with a luminance of 88.5 cd/m^2^. All Gabor patches were 2.75 degrees visual angle in diameter, presented at a 5 degree distance to the fixation dot (centre to centre) (all distances are shown to scale in **Supplementary Figure S1**). The fixation dot was 0.2 degrees in diameter and near-white (rgb: 234, 234, 234). The orientation change that occurred as the target event was always 2° clockwise or anticlockwise. Orientations of Gabor patches were randomly generated from a 160°-space, where orientations from -85 to -5 and 5 to 85 degrees were allowed, to avoid cardinal orientations.

### Experiment 1 vs. Experiment 2: Gaze-contingent trial termination

We conducted two complementary versions of the above described experiment that each employed a distinct but widely adopted approach to gaze control during the task. In Experiment 1, participants were instructed to fixate on the central fixation dot at all times and this was monitored by the experimenter who would repeat fixation instructions when necessary.

In contrast, during Experiment 2, trials were terminated when fixation strayed more than 2 degrees visual angle from the centre pixel in any direction. This set-up was attained by custom code that checked the current gaze position every 15 ms. Every gaze sample was checked in real-time and the trial was terminated as soon as any single sample was out of the allowed bounds. Upon trial termination, instructions to return to fixation were shown for 1 second. After this, the fixation dot was shown on its own for half a second, so participants could get their fixation back in order before the task continued with the next trial. Terminated trials were not replaced.

In Experiment 2, all trials where fixation was broken before the target event occurred were not included in both the behavioural and eye-tracking analyses. For the numbers of included trials and included saccades of both experiments, see **Supplementary Table S2**.

### Eye-tracking acquisition and preprocessing

Horizontal and vertical gaze positions of the participant’s right eye were continuously sampled using an EyeLink 1000 Plus at a rate of 1000 Hz throughout the experiment. The eye-tracker was positioned 10 cm in front of the monitor, and 60 cm away from the eyes. Participants used a chinrest to minimize head movements. Prior to recording, the eye tracker was calibrated and validated using the built-in HV9 calibration module. The eye tracker was always re-calibrated exactly halfway through the experiment (after 10 blocks), and could additionally be re-calibrated in each break, if the signal was deemed of too poor quality.

Following acquisition, eye-tracker datafiles were converted from their original Eyelink Data Format (.edf) to an ASCII text file (.asc), and analysed in MATLAB R2024b using a combination of the Fieldtrip analysis toolbox (47) and custom code. Custom code was used to detect blinks and remove them from the signal for 100 ms before and after each blink (by replacing these data with NaNs). After blink removal, data were epoched relative to cue onset. Finally, all trials with a premature keyboard response were removed from the saccade data analyses. In Experiment 2, we additionally removed all trials that were terminated by a break in fixation before the target event had occurred. In total, this yielded an average of 752±45 (M±SD, out of 800) trials per participant for Experiment 1 and 633±74 for Experiment 2.

For our main analysis (the results of which are shown in **Figure 2**) we exclusively analysed directional microsaccade biases prior to the target event (i.e., the orientation change of either visual stimulus). This ensures our results are uncontaminated from potential responses related to the target event itself. To also ensure sufficient trials for analysing microsaccades, we included all trials where the orientation change occurred later than 1400 ms after cue onset. This provided a good trade-off between having sufficient trials, while also having sufficient time to capture microsaccade biases during the initial shifting phase and the ensuing phase in which attention was maintained on the cued stimulus after this shift.

Importantly, our conclusions remain unaffected when investigating the entire time course through a complementary analysis (see **Supplementary Figure S2**). In this analysis, all data occurring after the orientation change were removed on each trial (by replacing these data with NaNs), thereby also ensuring our results are uncontaminated from potential responses related to the target event itself. While this complementary analysis does allow for examination of the full time course, the disadvantage of this approach is that the number of available data points decreases progressively towards the end. Consequently, statistical inferences become less stable at later time points. Because of this limitation, we did not adopt this approach as our main analysis.

### Saccade detection and visualisation

To detect saccades, a previously established and validated velocity-based method was employed (18) (building on (14)). In this approach, saccades are detected by finding those samples where the 2-dimensional velocity exceeds a certain trial-based threshold. Gaze velocity was obtained as the derivative of gaze position that was smoothed in the temporal dimension with a Gaussian-weighted moving mean filter with a 7-ms sliding window (using the built-in MATLAB function ‘smoothdata’). The velocity threshold was set at 5 times the median velocity in any given trial, which is consistent with earlier work (16, 19, 23). A minimum delay of 100 ms between successive saccades was imposed to avoid counting the same saccade multiple times. No criteria were set for minimum microsaccade duration or amplitude, but a visualisation of the relationship between saccade amplitude and peak velocity shows that the detected saccades generally follow the expected main sequence (see **Supplementary Figure S7**). Crucially, we did not restrict our analyses to initial saccades; rather, all detected saccades were included. This allowed us to examine oculomotor behaviour during later trial phases, where initial saccades are unlikely to occur.

Saccade size (in degrees) and direction were calculated by estimating the difference between the pre-saccade gaze position (−50 to 0 ms before threshold crossing) and the post-saccade gaze position (50 to 100 ms after threshold crossing). Because stimuli were placed at a 45° angle *below* the central fixation dot we only included downwards saccades (saccades with an angle between 0° and 180°; for a visualisation of which saccades were included, see **Supplementary Figure S3**), following the same logic as in (33). This ensured that all saccades were either toward the cued stimulus or toward the other stimulus, while neglecting upward saccades back to the fixation marker. (Note how this deliberate analysis choice also renders saccade rates lower than in many other articles that include saccades in all directions).

Depending on the direction of the saccade and the side of the cued item, we thus classified saccades as ‘towards the cued stimulus’ or ‘towards the other stimulus’, where the 180°-space was divided in two, so for a rightwards cue, all saccades between 0° and 90° were classified as ‘towards the cued stimulus’. Importantly, a complementary analysis shows our conclusions remain unaffected when the spatial saccade bias is investigated with narrower angular definitions of ‘towards cued stimulus’ and ‘towards other stimulus’ (see **Supplementary Figure S8**). After detecting and classifying the saccades, the participant’s individual time courses of saccade rates (in Hz) were determined by using a sliding time window of 100 ms, advancing in steps of 1 ms. These participant-level average time courses were smoothed using a Gaussian sliding time window of 200 ms, before subjecting these time courses to second level statistics using cluster-based permutation analysis.

To also investigate the size of the saccades that contributed to our findings without imposing an arbitrary threshold, we additionally decomposed saccade rates into a time-size representation, showing the time courses of saccade rates as a function of the saccade size (as also done in 16, 18, 23). For this, we used successive saccade-size bins of 0.5 degrees visual angle in steps of 0.1 degree.

### Statistical analysis

For the analysis of the behavioural data, we employed paired-sample Student’s t-tests to compare average reaction times and accuracy scores between trials in which the target event occurred in the cued (valid cue) or other (invalid cue) stimulus, collapsed over all cue-target intervals. Effect sizes for each comparison were quantified using Cohen’s d.

To statistically evaluate the time series gaze data, we employ a non-parametric cluster-based permutation approach (48). The permutation analysis was conducted using Fieldtrip with default clustering settings (grouping adjacent time points that were significant in a mass-univariate comparison). A permutation distribution of the largest cluster size was acquired by randomly permuting the condition labels of each participant’s trial-averaged time course data (i.e. randomly flipping the sign of the difference in rate of toward vs. away saccades) 10,000 times and identifying the size of the largest clusters observed in these randomised data after each permutation. The p-values of the clusters observed in the original data were calculated as the proportion of random permutations where the largest cluster was equal to or larger than the cluster that we observed in the original data.

The cluster-based permutation approach only identifies temporal clusters of significance, but is not suited to directly compare modulations between the pre-defined time windows of shifting vs. maintaining attention. In order to compare the magnitude of the saccade bias between the two time windows of interest, we defined the shift window as the window between 200-600 ms after cue onset (determined a-priori, based on prior work, see 16, 18, 23 and consistent with ample related literature, see 35–37) and we defined the subsequent stage of attentional maintenance in the period following this (in the main analysis, this period lasts from 600 until 1400 ms after cue onset). This enabled us to average the saccade biases (defined as the difference in saccades rates toward the cued versus the other stimulus) in both a-priori defined time windows and to compare these biases using paired-sample Student’s t-tests. Effect sizes for each comparison were again quantified using Cohen’s d. As this analysis resulted in a non-significant p-value, we supplemented this analysis with a Bayesian analysis to also quantify evidence in favour of the absence of an effect. This was implemented in JASP using default settings (i.e., the prior distribution is a Cauchy distribution with location 0 and scale 0.707).

## Acknowledgements

This work was supported by an ERC Starting Grant from the European Research Council (MEMTICIPATION, 850636) and an NWO Vidi Grant from the Dutch Research Council (14721) to F.v.E.

## Data, Materials, and Software Availability

All study data and analysis code will be made public prior to publication. The code for the experimental task is already publicly available on [GitHub](https://github.com/annavanharmelen/Microsaccade-bias-experiment).

## Supplementary Information

**Table S1.** Cue-validity effects at each of the tested cue-target intervals. This table splits the cue-validity effects from Figure 1 into the ten possible cue-target intervals. The top half of the table shows the effects of cue-validity on reaction time; the bottom half of the table shows the effects of cue-validity on accuracy. Results from Experiment 1 and 2 are shown, respectively, on the left and right sides of the table. The table shows both uncorrected and Bonferroni corrected p-values, along with their significance levels: *: p< 0.05, **: p < 0.01, ***: p < 0.001.

| Reaction time |  |  |  |  |  |  |
| --- | --- | --- | --- | --- | --- | --- |
|  | Experiment 1 |  |  | Experiment 2 |  |  |
| Cue-target interval (ms) | t-value | p-value (uncorrected) | p-value (Bonferroni) | t-value | p-value (uncorrected) | p-value (Bonferroni) |
| 500 | 3.431 | 0.002** | 0.02* | 3.876 | 7.66e-04*** | 0.008** |
| 800 | 3.482 | 0.002** | 0.02* | 3.211 | 0.004** | 0.04* |
| 1100 | 3.007 | 0.006** | 0.06<sup>n.s.</sup> | 3.822 | 8.74e-04*** | 0.009** |
| 1400 | 4.022 | 5.33e-04*** | 0.005** | 3.864 | 7.88e-04*** | 0.008** |
| 1700 | 3.299 | 0.003** | 0.03* | 3.384 | 0.003** | 0.03* |
| 2000 | 3.322 | 0.003** | 0.03* | 2.717 | 0.012* | 0.12<sup>n.s.</sup> |
| 2300 | 3.278 | 0.003** | 0.03* | 3.347 | 0.003** | 0.03* |
| 2600 | 2.604 | 0.016* | 0.16<sup>n.s.</sup> | 2.826 | 0.010* | 0.1<sup>n.s.</sup> |
| 2900 | 3.629 | 0.001** | 0.01* | 3.422 | 0.002** | 0.02* |
| 3200 | 1.603 | 0.122<sup>n.s.</sup> | 1.00<sup>n.s.</sup> | 3.390 | 0.003** | 0.03* |
| Accuracy |  |  |  |  |  |  |
|  | Experiment 1 |  |  | Experiment 2 |  |  |
| Cue-target interval (ms) | t-value | p-value (uncorrected) | p-value (Bonferroni) | t-value | p-value (uncorrected) | p-value (Bonferroni) |
| 500 | 2.389 | 0.025* | 0.25<sup>n.s.</sup> | 3.505 | 0.002** | 0.02* |
| 800 | 2.322 | 0.029* | 0.29<sup>n.s.</sup> | 2.713 | 0.012* | 0.12<sup>n.s.</sup> |
| 1100 | 2.934 | 0.007* | 0.07<sup>n.s.</sup> | 3.488 | 0.002** | 0.02* |
| 1400 | 2.310 | 0.030* | 0.3<sup>n.s.</sup> | 3.190 | 0.004** | 0.04* |
| 1700 | 1.893 | 0.071<sup>n.s.</sup> | 0.71<sup>n.s.</sup> | 2.356 | 0.027* | 0.27<sup>n.s.</sup> |
| 2000 | 1.363 | 0.186<sup>n.s.</sup> | 1.00<sup>n.s.</sup> | 1.728 | 0.097<sup>n.s.</sup> | 0.97<sup>n.s.</sup> |
| 2300 | 1.976 | 0.060<sup>n.s.</sup> | 0.6<sup>n.s.</sup> | 2.382 | 0.026* | 0.26<sup>n.s.</sup> |
| 2600 | 1.399 | 0.175<sup>n.s.</sup> | 1.00<sup>n.s.</sup> | 1.804 | 0.084<sup>n.s.</sup> | 0.84<sup>n.s.</sup> |
| 2900 | 2.419 | 0.024* | 0.24<sup>n.s.</sup> | 1.303 | 0.205<sup>n.s.</sup> | 1.00<sup>n.s.</sup> |
| 3200 | 1.783 | 0.088* | 0.88<sup>n.s.</sup> | 2.039 | 0.053<sup>n.s.</sup> | 0.53<sup>n.s.</sup> |

**Table S2.** Average number of trials per participant per condition and average number of saccades per participant per condition. This table reflects the amount of data included in the main analyses shown in Figure 1, Figure 2 and Figure 3.

|  | Trials included behaviourally per subject per condition (M±SD) |  | Saccades per subject per condition (M±SD) |  |  |  |
| --- | --- | --- | --- | --- | --- | --- |
|  |  |  | Shift |  | Maintain |  |
|  | Valid | Invalid | Toward | Away | Toward | Away |
| **Experiment 1** | 640±0 | 160±0 | 39±30 | 24±18 | 42±39 | 27±22 |
| **Experiment 2** | 529±54 | 133±14 | 18±18 | 16±16 | 19±20 | 18±21 |

**Figure S1.**
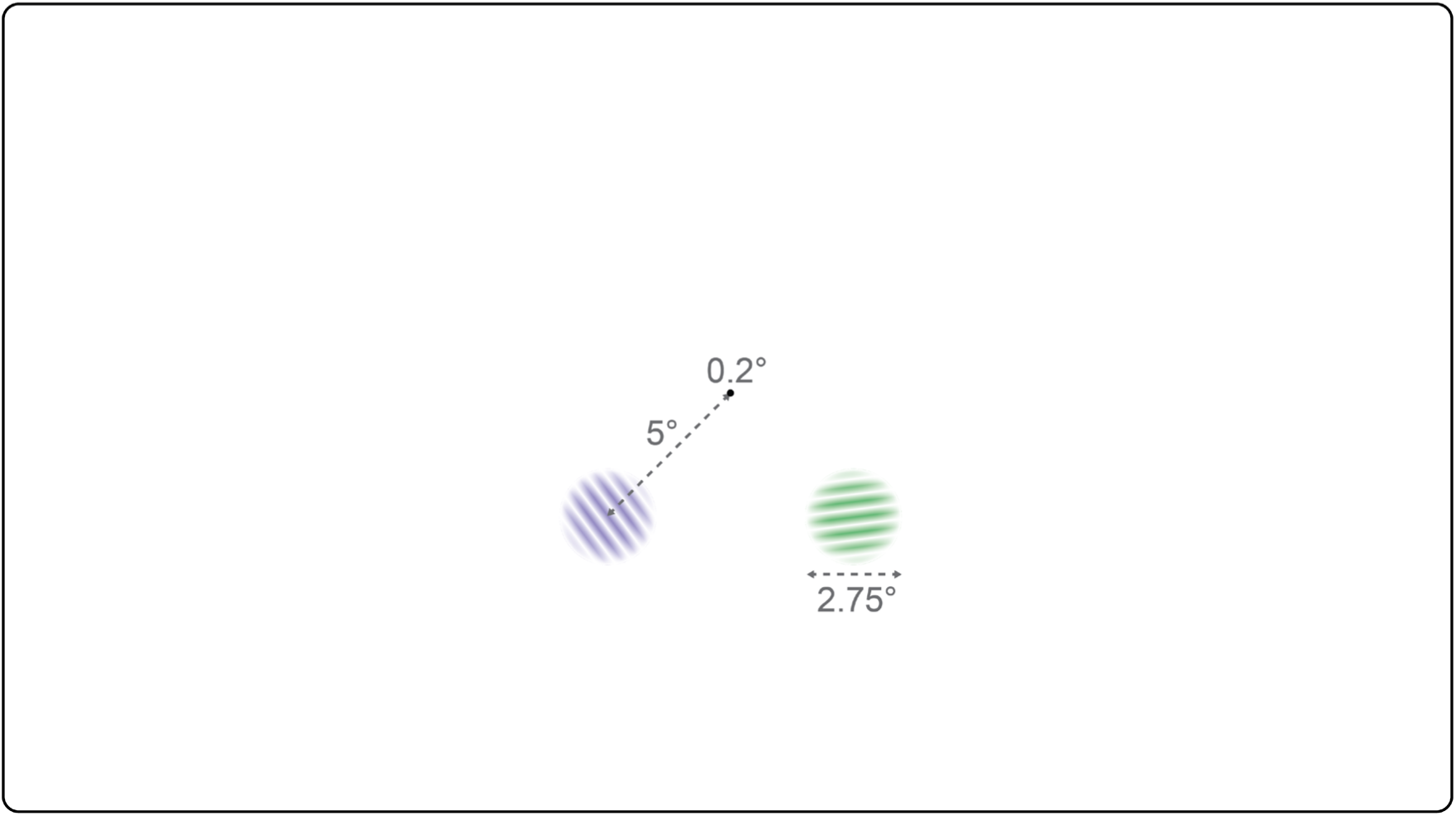
Size of the experimental screen, stimuli, and distances to scale.

Attention was successfully shifted and maintained throughout the entire trial period.

**Figure S2.**
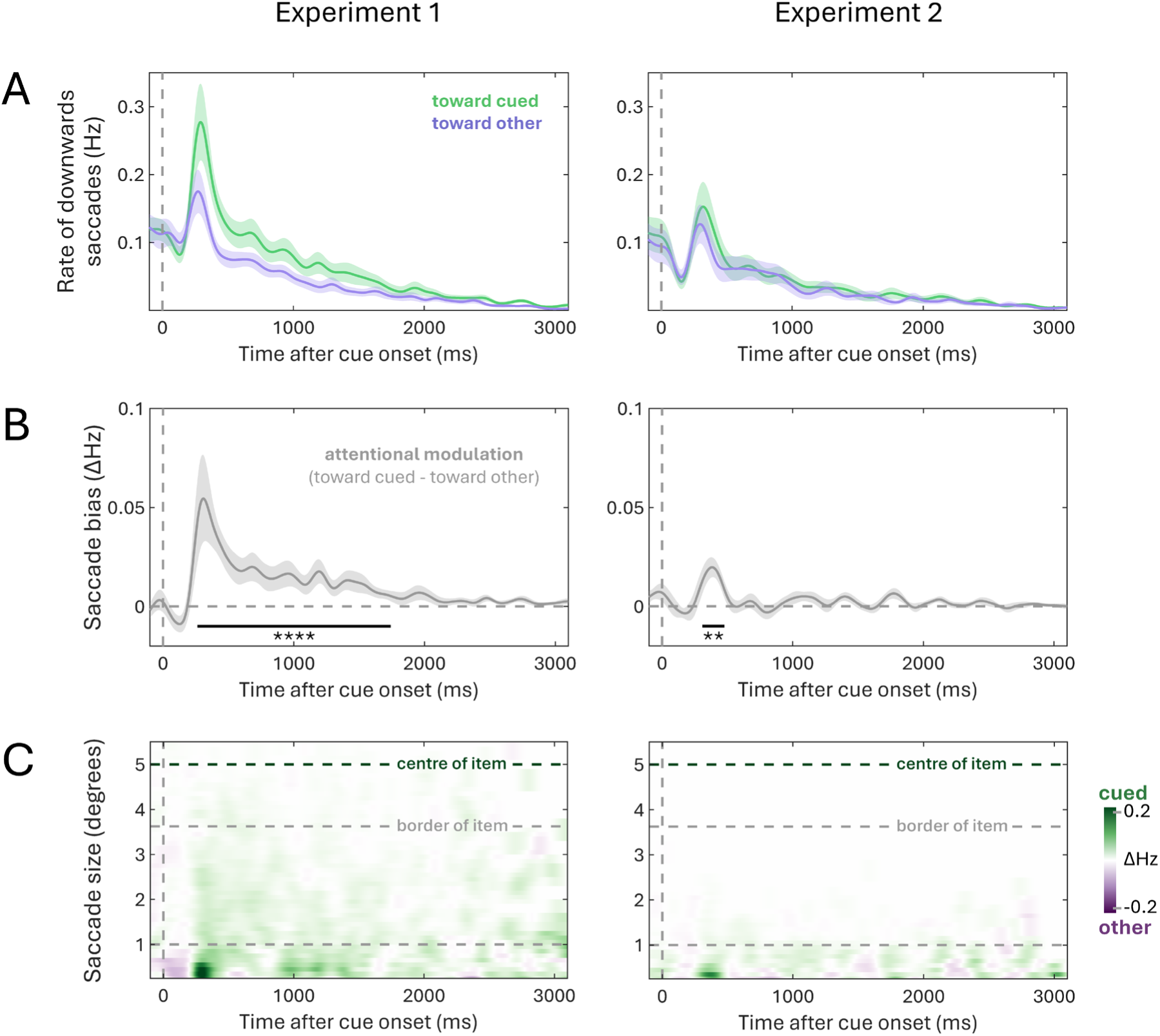
Full time courses of saccade data. This figure replicates the results shown in Figure 2 using the complementary analysis described in the Methods section. (A) Time courses of saccade rates for saccades towards the cued and the other item. This analysis only includes downward saccades, as stimuli were positioned at a 45° angle below fixation, allowing us to distinguish saccades toward the item from those returning to central fixation. The time courses indicate mean values, with shaded areas indicating the standard error of the mean. (B) Time courses of the attentional modulation of saccade direction (toward cued – toward other). The time courses indicate mean values, with shaded areas indicating the standard error of the mean. Black horizontal lines indicate significant temporal clusters (as determined by a cluster-based permutation test). (C) Time courses of attentional modulation (as shown in B) as a function of saccade size. For reference, dashed horizontal lines indicate 1 degree visual angle, as well as the locations of the centre and closest border of the visual stimulus. Throughout the entire figure, the following significance levels were used: *: p< 0.05, **: p < 0.01, ***: p < 0.001, ****: p < 0.0001.

**Figure S3.**
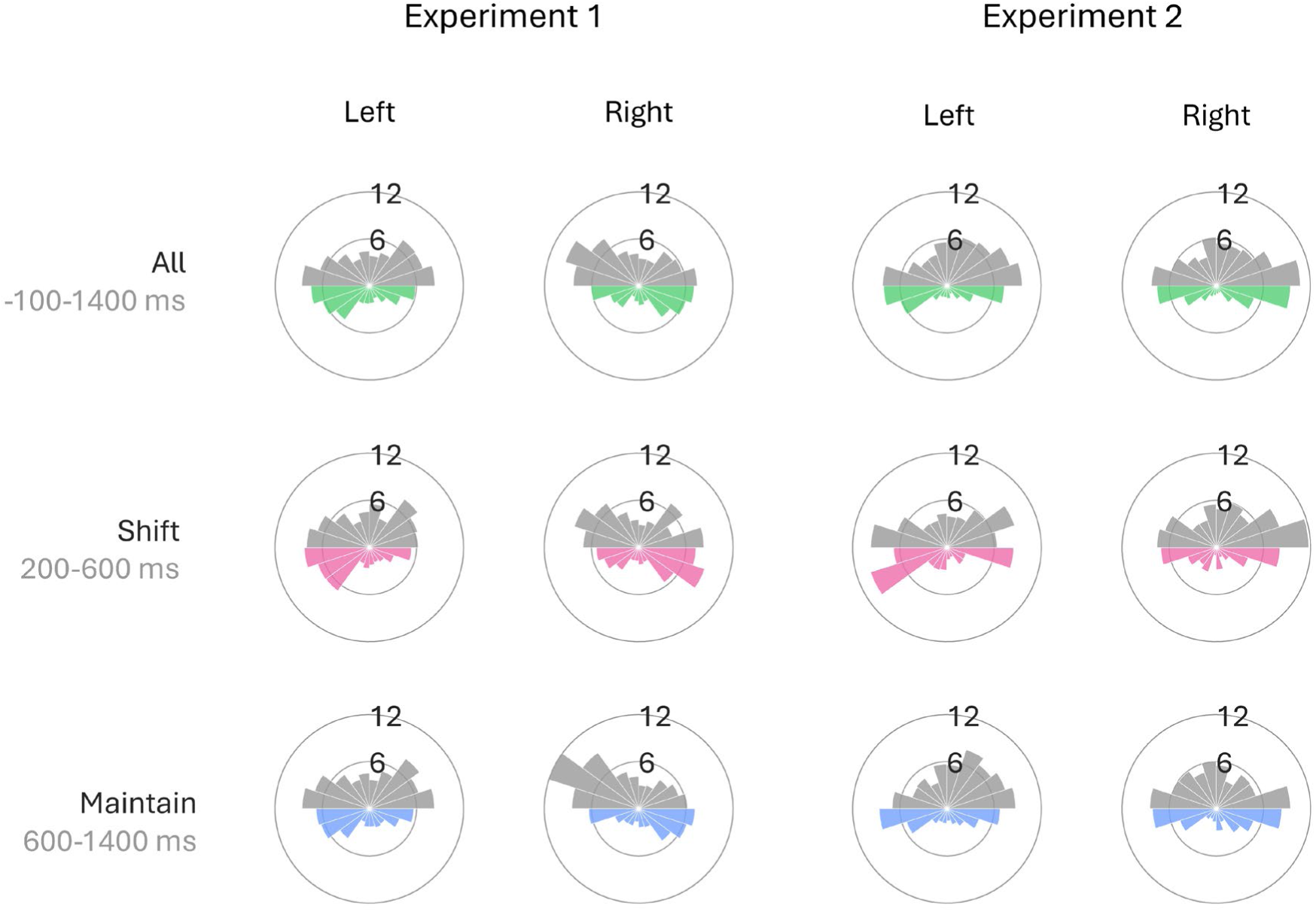
Polar histograms of all saccades included in the main analyses. This figure shows the percentage of detected saccades going in a specific direction for both experiments. This visualisation is repeated for the three main timeframes of interest: (1) the whole trial (from -100 to 1400 ms after cue onset), (2) the ‘shift’ period (from 200 to 600 ms after cue onset) and (3) the ‘maintain’ period (from 600 to 1400 ms after cue onset). The coloured saccades are those classified as ‘downwards’ saccades in the analyses of Figure 2. The radial axis shows the percentage of detected saccades going in one of twenty binned possible directions, which is calculated by weighting each participant equally (i.e., averaging the proportion of saccades detected for each participant per polar angle bin).

**Figure S4.**
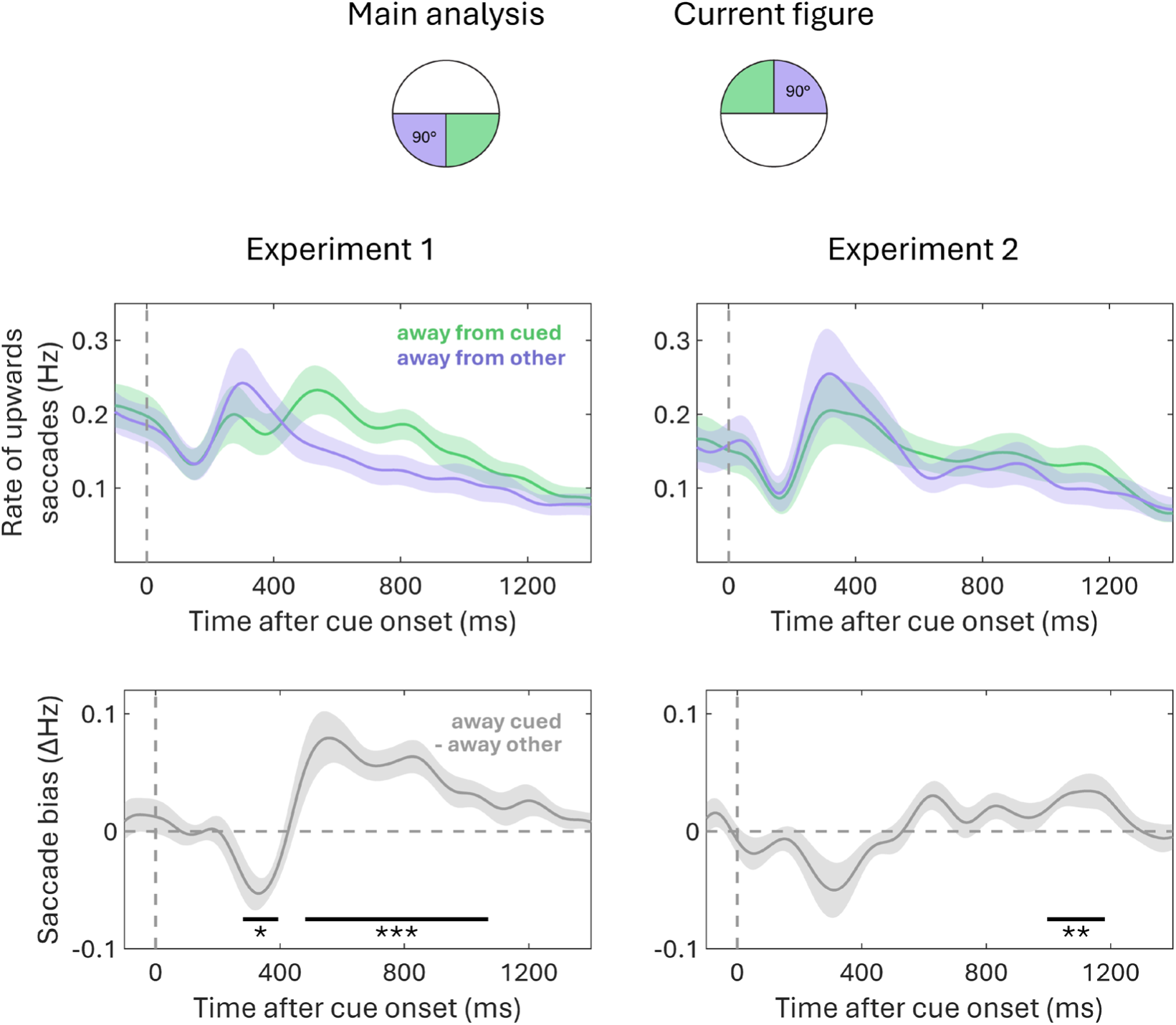
Attentional bias in upwards saccades. This figure repeats the analysis shown in Figure 2, but for upwards saccades instead of downwards saccades. Note that the visual items were always downward in our task. (A) Time courses of saccade rates for saccades away from the cued and the other item. This analysis only includes upwards saccades. As stimuli were positioned at a 45° angle below fixation, this mainly reflects a return to central fixation. The time courses indicate mean values, with shaded areas indicating the standard error of the mean. (B) Time courses of the attentional modulation of saccade direction (away cued – away other). The time courses indicate mean values, with shaded areas indicating the standard error of the mean. Black horizontal lines indicate significant temporal clusters (as determined by a cluster-based permutation test). Throughout the entire figure, the following significance levels were used: *: p< 0.05, **: p < 0.01, ***: p < 0.001, ****: p < 0.0001.

In Experiment 1 a positive correlation was found between the magnitude of the cue-related attentional modulation and the magnitude of the spatial saccade bias, but this did not replicate in Experiment 2.

**Figure S5.**
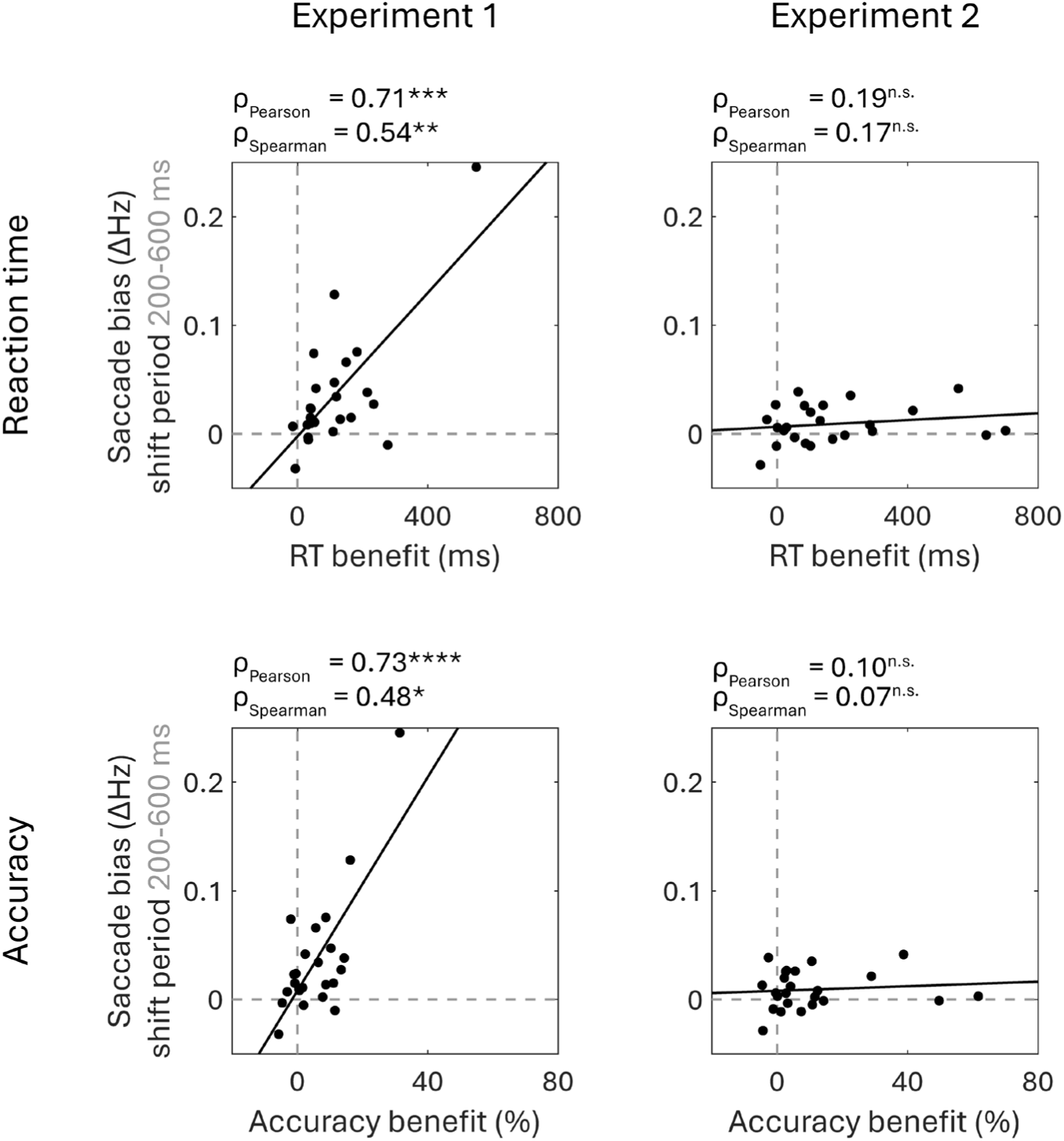
Relationship between the magnitude of the cue-related attentional modulation and the magnitude of the spatial saccade bias. This figure shows the relationship between the cue-related attentional modulation (referred to in the figure as a ‘benefit’ for simplicity) and the average saccade bias during the ‘shift’ period (from 200 to 600 ms after cue onset), for both reaction time and accuracy. Each dot represents one participant. Throughout the entire figure, the following significance levels were used: *: p< 0.05, **: p < 0.01, ***: p < 0.001, ****: p < 0.0001.

Fixational control was successful in both experiments.

**Figure S6.**
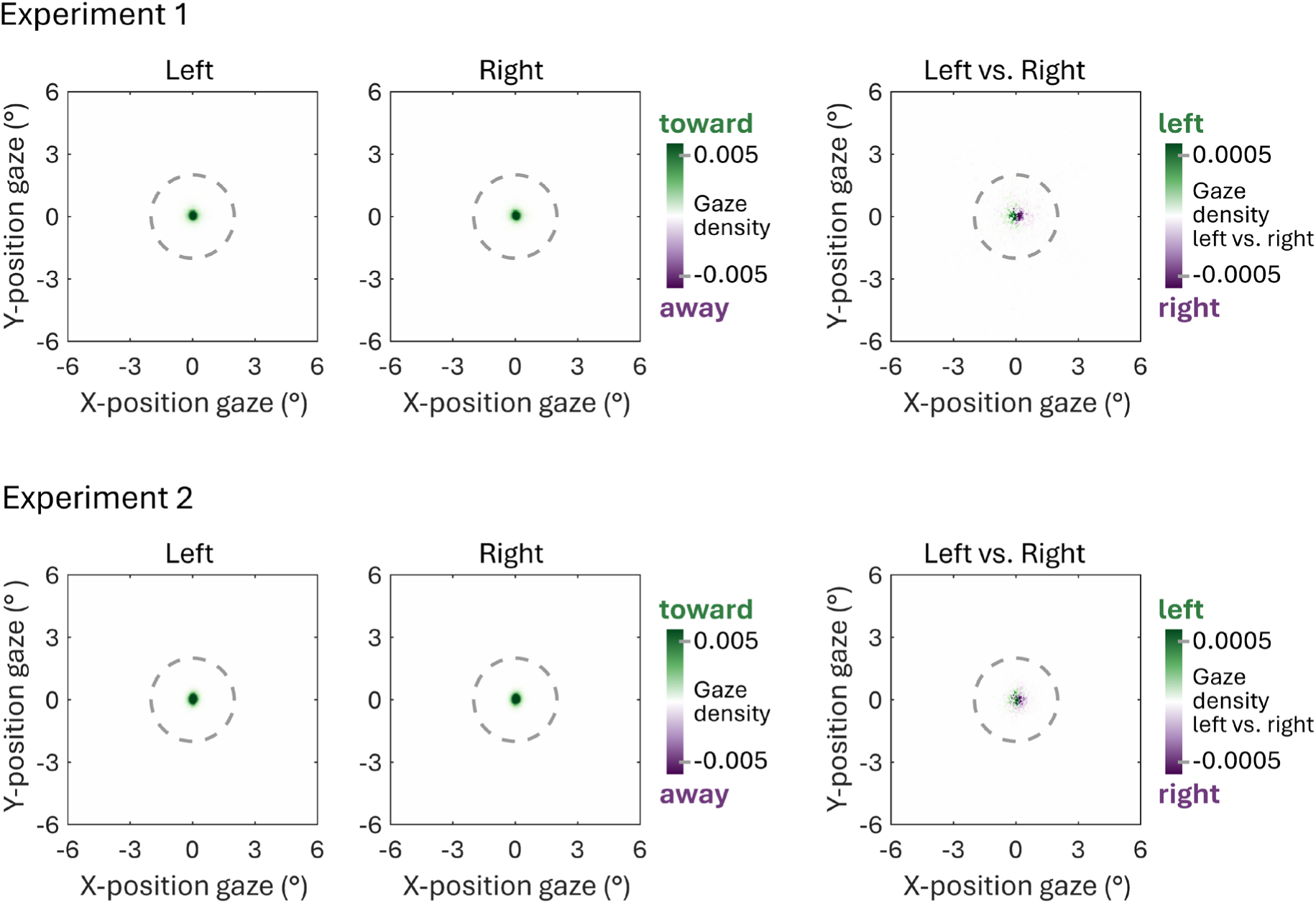
Gaze density during the ‘shift’ period. This figure shows the gaze density during the ‘shift’ period (from 200 to 600 ms after cue onset) for both experiments, separately for left cued trials and right cued trials. The very right column shows the difference in gaze density between left and right cued trials.

In both experiments, the saccade-detection algorithm worked well: detected saccades generally follow the main sequence.

**Figure S7.**
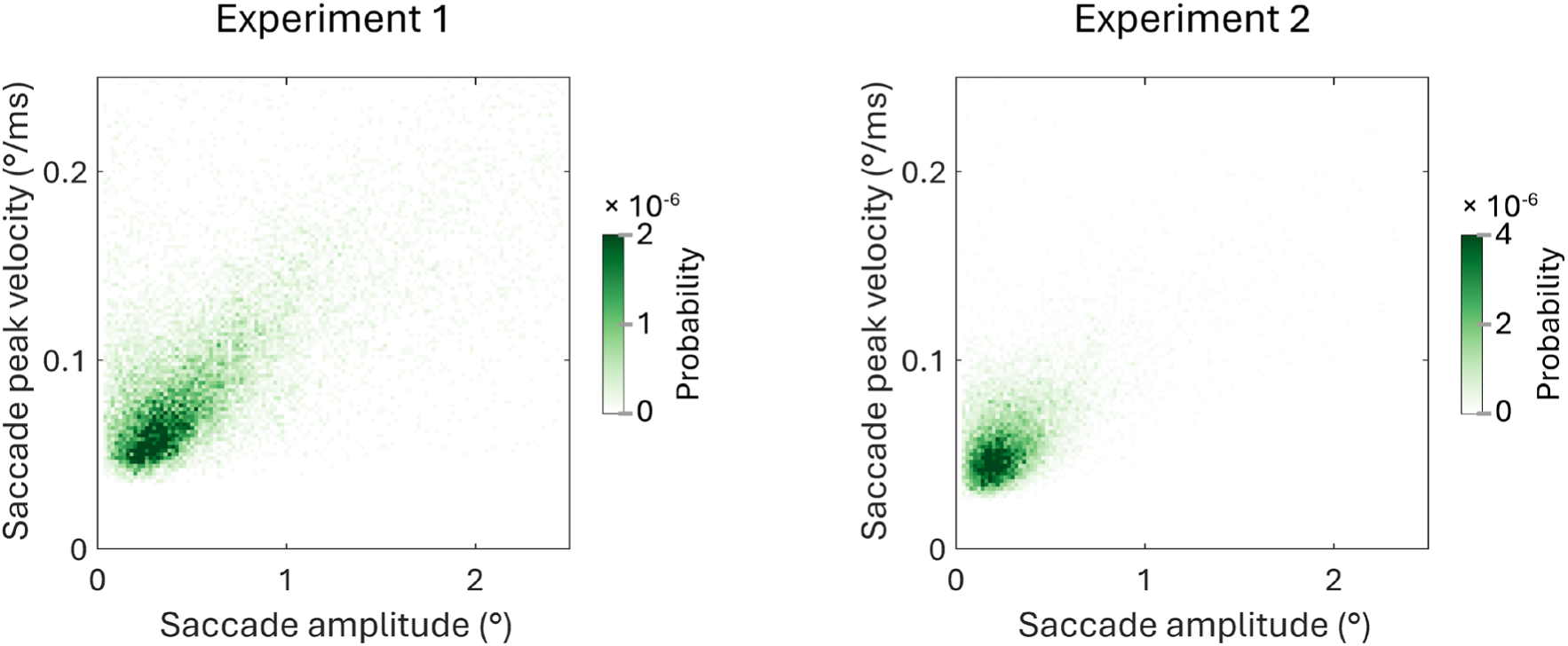
Relationship between amplitude and peak velocity of all detected saccades. This figure shows the relationship between the amplitude (in degrees visual angle) and peak velocity (in degrees visual angle / millisecond), for Experiment 1 and Experiment 2 separately. A visual inspection of both plots shows that the detected saccade generally follow the main sequence.

The main results of both experiments are replicated when using a narrower angular definition of ‘toward cued’ and ‘toward other’.

**Figure S8.**
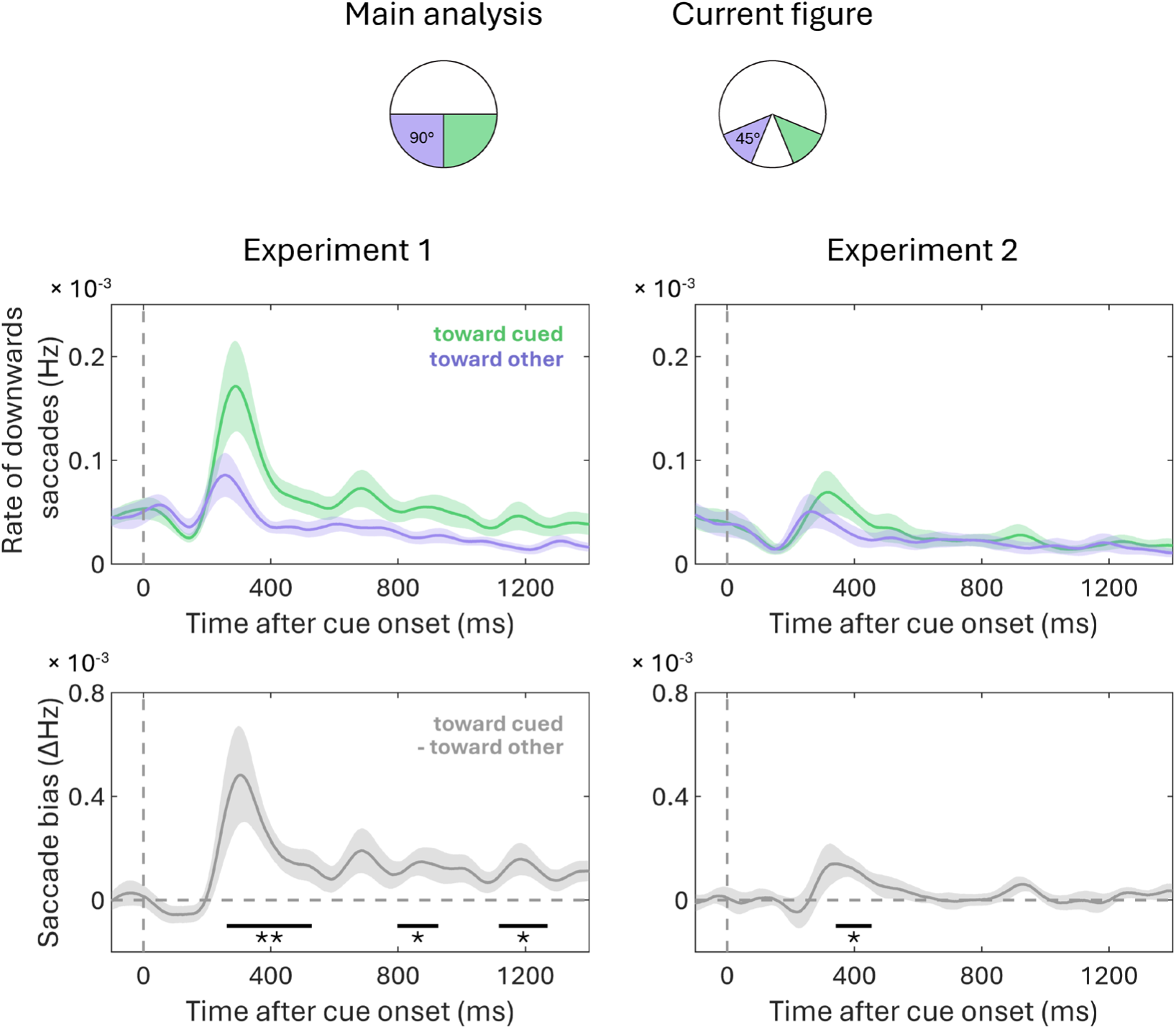
Re-analysis of saccade data with narrower angular definitions of ‘toward cued’ and ‘toward other’. This figure repeats the analysis shown in Figure 2, but uses an angular definition of 45° around the direction of the visual stimulus to define the ‘toward cued’ and ‘toward other’ directions, instead of the original 90°. (A) Time courses of saccade rates for saccades towards the cued and the other item. This analysis only includes downward saccades, as stimuli were positioned at a 45° angle below fixation, allowing us to distinguish saccades towards the item from those returning to central fixation. The time courses indicate mean values, with shaded areas indicating the standard error of the mean. (B) Time courses of the attentional modulation of saccade direction (toward cued – toward other). The time courses indicate mean values, with shaded areas indicating the standard error of the mean. Black horizontal lines indicate significant temporal clusters (as determined by a cluster-based permutation test). Throughout the entire figure, the following significance levels were used: *: p< 0.05, **: p < 0.01, ***: p < 0.001, ****: p < 0.0001.

